# Exploring bycatch diversity of organisms in whole genome sequencing of Erebidae moths (Lepidoptera)

**DOI:** 10.1101/2021.09.02.458197

**Authors:** Hamid Reza Ghanavi, Victoria Twort, Anne Duplouy

## Abstract

Models estimate that up to 80% of all butterfly and moth species host vertically transmitted endosymbiotic microorganisms, which can affect the host fitness, metabolism, reproduction, population dynamics, and genetic diversity, among others. The supporting empirical data are however currently highly biased towards the generally more colourful butterflies, and include less information about moths. Additionally, studies of symbiotic partners of Lepidoptera predominantly focus on the common bacterium *Wolbachia pipientis*, while infections by other inherited microbial partners have more rarely been investigated. Here, we mine the whole genome sequence data of 47 species of Erebidae moths, with the aims to both inform on the diversity of symbionts potentially associated with this Lepidoptera group, and discuss the potential of metagenomic approaches to inform on their associated microbiome diversity. Based on the result of Kraken2 and MetaPhlAn2 analyses, we found clear evidence of the presence of *Wolbachia* in four species. Our result also suggests the presence of three other bacterial symbionts (*Burkholderia* spp., *Sodalis* spp. and *Arsenophonus* spp.), in three other moth species. Additionally, we recovered genomic material from bracovirus in about half of our samples. The detection of the latter, usually found in mutualistic association to braconid parasitoid wasps, may inform on host-parasite interactions that take place in the natural habitat of the Erebidae moths, suggesting either contamination with material from species of the host community network, or horizontal transfer of members of the microbiome between interacting species.

## Introduction

A growing scientific community now sees each organism as a community of interacting species rather than as an independent entity. Insects are no exception. They host a variety of microbial symbionts sitting both inside and outside their host cells. These microorganisms are at least as numerous as the number of host cells, and may constitute up to 10% of the host total mass ^1^. The effects of symbionts on their insect hosts are potentially as diverse as their taxonomy, ranging from pathogenic to obligate mutualists, and all the intermediate possible relationships ^2^. This diversity has recently attracted the growing interest of the scientific community, but gaps and biases remain. For example, in Lepidoptera, research in symbiosis has mostly focused on the most charismatic groups of colourful diurnal butterflies ^3–5^ and on pest species to the human society ^6–8^. In contrast, the rest of Lepidoptera (mainly moths) which encompass no less than 130,000 species ^9^, have rarely been screened for their associations with symbionts ^10^.

High throughput sequencing technologies (HTS) now provide a relatively easy and cheap way to obtain large amounts of genetic data. These technologies used to generate genomic data are varied and broadly applicable to the widest range of organisms. Thereby, revolutionizing our accessibility to genomic resources and continually expanding and renewing the scope of the questions we can address within the natural sciences. For example, sequencing material from a particular study organism, either entirely or partially, may results in a mix of primary host specific DNA and DNA from other sources. These other sources can include ecto/endosymbionts, food, and opportunistic parasites and pathogens, among others. Such genomic data opens up the genomic analyses towards broader targets, especially towards investigating the diversity of symbionts that might be associated to particular targeted hosts. Here, we mine the data produced from whole genome sequencing of 47 moth species from the family Erebidae to i) explore the potential diversity of symbionts associated to this megadiverse Lepidoptera family; and ii) to evaluate the exploratory power of recovering information on natural host-symbiont associations from the low coverage genome sequencing approaches.

## Results

### Metagenomic analysis

We identified the species *Idia aemula, Luceria striata, Acantholipes circumdata* and *Oraesia excavata* (RZ271, RZ42, RZ248, and RZ337) as infected by *Wolbachia*, and *Wolbachia*-associated phage *WO* (Table 1), with between 66,978 and 208,044 of the reads identified as belonging to the symbiont. Additionally, the reads obtained from sample RZ13 (*Gonitis involuta*) was also found to include 954 *Wolbachia* reads, which is a higher number of reads than found for any of the clearly uninfected specimens, but is considerably less than any of the four clearly infected specimens listed above.

**Table 1.**
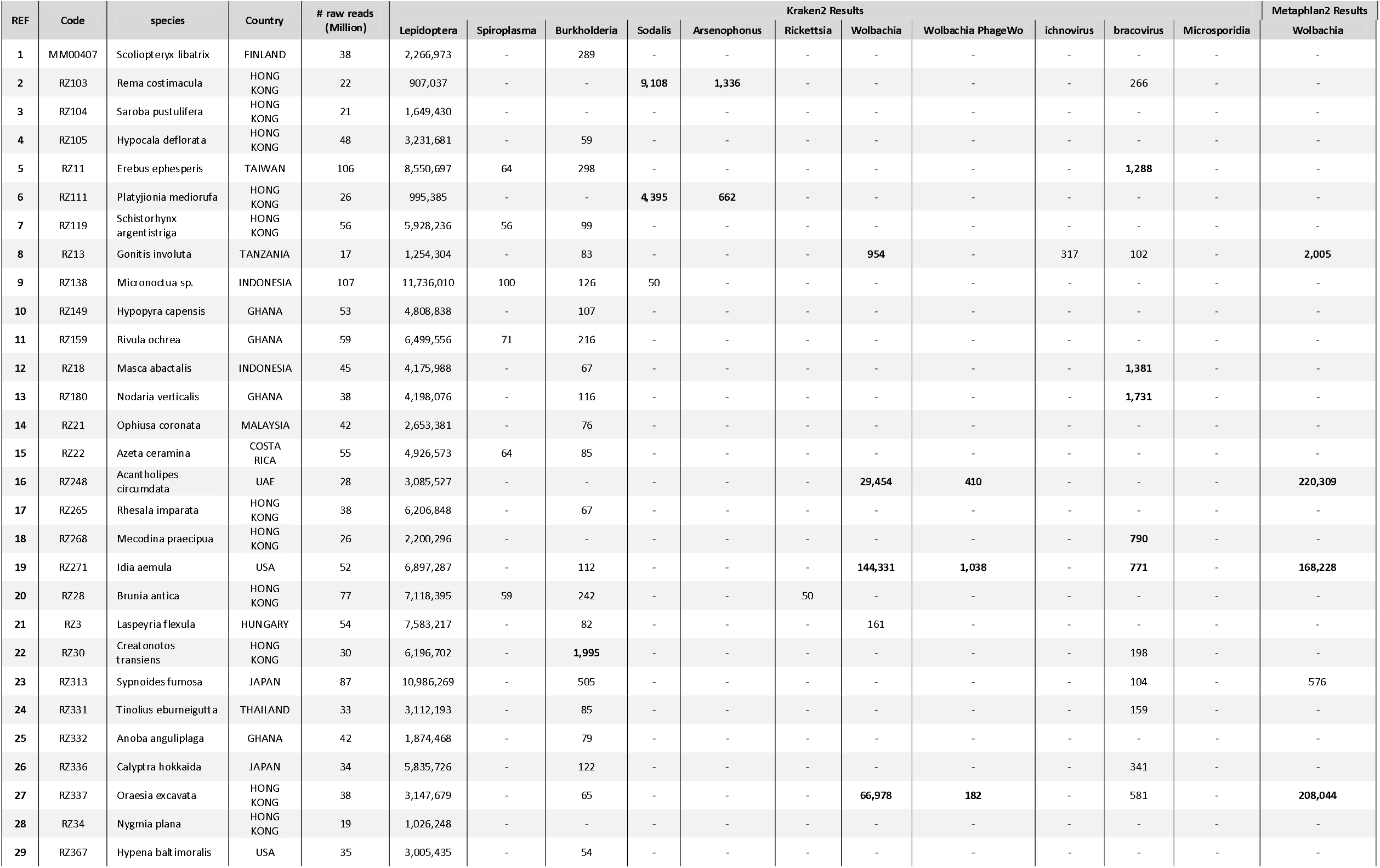
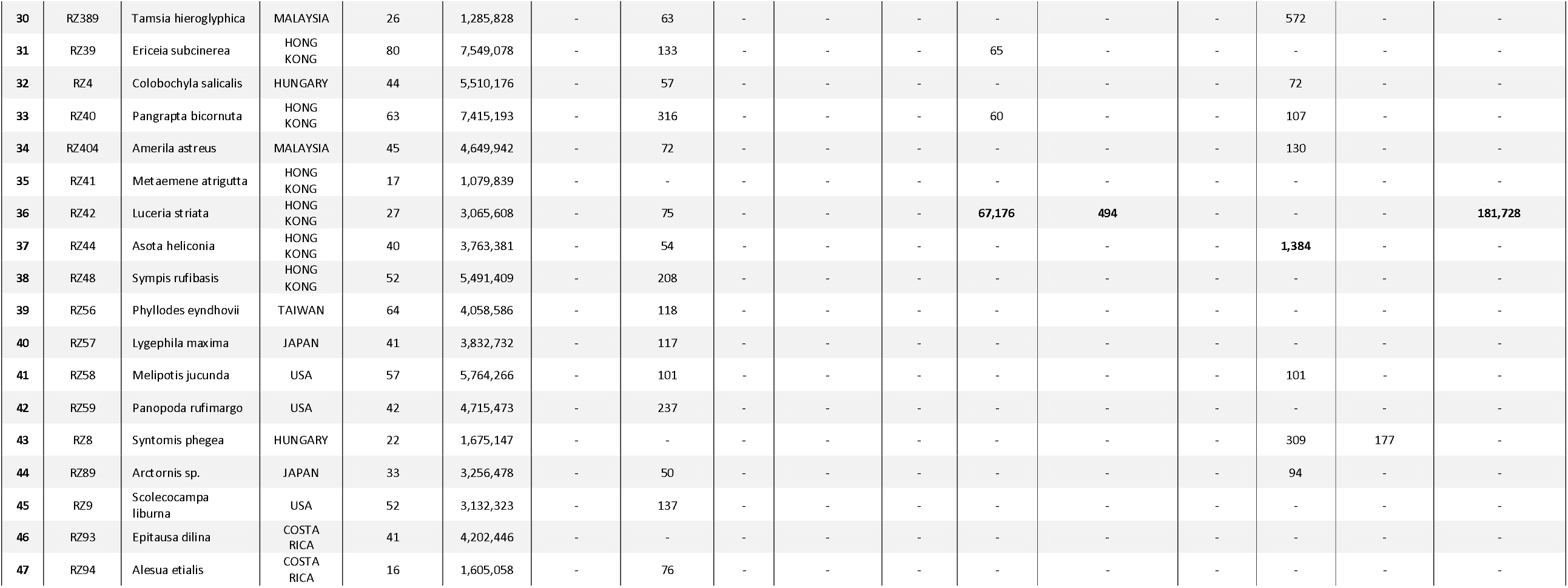
The number of reads classified as originating from the host and various microorganisms. Values in bold highlight the values mentioned in the text, - represent samples with either zero or less than 50 reads classified.

Our Kraken2 and MetaPhlan2 analyses showed no to very few reads mapping to *Cardinium, Hamiltonella* or *Spiroplasma* bacteria, or to Microsporidian fungi, in any of the 47 datasets screened. In contrast, the specimens RZ103 and RZ111 (*Rema costimacula* and *Platyjionia mediorufa*) included considerably more reads from *Sodalis* bacteria (9,108 and 4,395, respectively), and from *Arsenophonus* bacteria (1,336 and 662, respectively), than any other samples (maximum of 50 reads in any other sample). A closer look at the Kraken2 outputs from the latter two samples also revealed a possible infection with a *Plautia stali* symbiont (gammaproteobacteria; 3,856 and 1,914 reads, respectively), which was not detected in any of the other 45 samples. Additionally, the sample RZ30 (*Creatonotos transiens*) is the only one to show relatively high number of reads mapping to *Burkholderia* bacteria (N=1,995). Finally, we identified a considerable amount of reads from viruses of the polydnaviridae family, and especially of the Bracoviruses in three samples, *Erebus ephesperis, Masca abactalis* and *Asota heliconia* (RZ11, 1288 reads, RZ18, 1381 reads, and RZ44, 1384 reads). All other samples only include less than 750 reads, and more often no reads, for these viruses.

All details of the screen for the common symbionts can be found in Table 1, while all results from the Kraken2 and MetaPhlAn2 analyses can be found in the supplementary material and GitHub repository.

## Discussion

We confidently add four moth species (i.e., *Idia aemula, Luceria striata, Acantholipes circumdata* and *Oraesia excavata*) to the list of species hosting the intracellular alpha-proteobacterial symbiont *Wolbachia* (Hornett and Duplouy 2018), confirmed through two screening methods (i.e., Kraken2 and MetaPhlAn). With only 4 out of 47 species (8%) found infected, this represents a lower infection rate than presented in the current literature suggests (i.e., 16-79% of the studied lepidopteran groups infected with *Wolbachia*; ^11–16^. The general penetrance of *Wolbachia* however varies significantly among species, and is often low within infected populations ^17^. Thus, our results are most likely underestimating the true infection rate within the Erebidae moths. Future broader screenings of different populations will provide more accurate natural infection rates for these species.

Noticeably, we observe the presence of *Wolbachia* phage *WO* within those samples for which *Wolbachia* presence is strongly supported. The interaction of this bacteriophage with *Wolbachia* has been the focus of many evolutionary studies in recent years ^18–22^. Previous research suggests that phage *WO* are associated with horizontal gene transfer in *Wolbachia*, and with genes that may affect the fitness of the bacterium ^23,24^. These bacteriophages have been observed in practically all the studied genomes of *Wolbachia* up to date, with very few obligate mutualistic exceptions ^18,25,26^. In the sample RZ13, species *Gonitis involuta*, a relatively high number of reads mapped to *Wolbachia* (1K reads), although significantly lower than in the other four species (29K-144K reads), and no reads were mapped to phage-*WO*. Few non-excluding hypotheses may explain such a pattern, these reads might originate from (I) contamination with other genetic material alien to our sample, (II) the integration of *Wolbachia* genomic material (partially or entirely) in the host genome, (III) random errors in the Identification of the reads as *Wolbachia*, (IV) low quality genomic material or (V) a combination of above-mentioned reasons. The overall screening results suggest that this sample was of low quality prior to sequencing. We however cannot rule out any of the other possibilities, and more studies are needed to fully confirm or reject the presence of *Wolbachia* in this species.

The two moth samples, *Rema costimacula* (RZ103) and *Platyjionia mediorufa* (RZ111), were of particular interests. Both the Kraken2 and the MetaPhlAn2 analyses suggest the presence of three gammaproteobacteria endosymbionts, namely *Sodalis, Arsenophonus and* ‘*Plautia stali-*symbiont’ in both samples. *Sodalis* has been characterized from different insects, including tsetse flies ^27^, seal louse ^28^, pigeon louse ^29^, loose flies ^30^, aphids ^31^, seed bug ^32^, weevils ^33,34^, stinkbugs ^35^, bees ^36^, and ants ^37^, among others. To our best knowledge however, this is the first time the three symbionts are found in Lepidoptera (Duplouy and Hornett 2018). This suggests that *Sodalis* bacteria might affect a more diverse group of organisms than is currently known. We are however cautious with the interpretation of this result, as the simple discovery of bacteria in the genomic data does not inform us about the nature of their interactions with the hosts. Whether *Sodalis* and the moth species share a symbiotic relationship, or not, will only be confirmed via experimentation and testing of the partnership through the host generations. Contamination of those two samples prior to DNA extraction is always possible. However, the sequenced host genetic material did not include significant amount of hemipteran DNA (or any other non-lepidopteran insect order), with comparable low numbers of reads (<1,500) mapped to Hemipterans in all the sequenced genomes. This, rules out DNA contamination by material from the previously confirmed hemipteran hosts of these three symbionts. It is shown that the female brown-winged green bug, *P. stali*, smears excrement over the egg surface during oviposition. The nymphs acquire the symbionts right after hatching by ingesting the excrements ^38^. Therefore, a possible contamination source could be any contact with such excrement/egg clusters. Once again, studies of the symbionts in natural populations of these moth species are needed to fully resolve the true infection state of these species and the relationship with the bacteria.

The moth species *Creatonotos transiens* shows a potential partnership with proteobacteria *Burkholderia* sp. In Lepidoptera, *Burkholderia* are known from the microbiota associated with the moth *Lymantria dispar* ^39^. However, similarly to the other symbionts presented above, these bacteria are also found in very diverse groups of organisms, from Amoebas to Orthoptera, from humans to plants ^40–43^. In the bean bug, *Riptortus pedestris*, studies have suggested that the bacteria can benefit their host by providing resistance to pesticides ^44^. Although never tested, the presence of such Proteobacteria in moths could similarly enhance the host ability to resist pesticides. If proven true, this could contribute to partially explaining the global success of many pest moth species despite the development of various targeted control strategies.

Six genomes included significantly high amounts of bracovirus reads, *Erebus ephesperis* (RZ11), *Masca abactalis* (RZ18), *Nodaria verticalis* (RZ180), *Mecodina praecipua* (RZ268), *Idia aemula* (RZ271) and *Asota heliconia* (RZ44). Bracoviruses are a known genus of mutualistic viruses with a complex life cycle. Integrated in the genome of a braconid parasitic wasp, the bracovirus is transcribed during oviposition in lepidopteran larvae ^45^. The presence of this viral genetic material in adult moths might suggest an unsuccessful infection by the parasitoid, and the survival of the larvae carrying the parasitic viral particles. Another potential explanation includes the possibility for the viral DNA to be integrated into the lepidopteran genome, as it is usually found in its common Hymenoptera host. Only studies simultaneously investigating parasitism success rate and tissue tropism of the bracoviruses in the Lepidoptera and Hymenoptera hosts, will be able to inform on the nature of these interactions.

From a methodological point of view, the present study shows the successful exploratory approach to mine for potentially hidden associated microbial diversity in genomic data. Our study was performed on shallow genome short reads obtained using Illumina platform. The original purpose of this sequencing effort was to study the phylogenomics of the hosts species ^46^, but a similar approach to the one we have taken here can be implemented to any publicly available genomic datasets. The popularity of genomic scale sequence data methods, such as Illumina short read approach, created a wide publicly open genomic resource for the research community to study questions that are not directly into the focus of the studies generating them. It is however important to also consider the limitations of such approaches. First, the quality and completeness of the reference datasets needed for programs like Kraken2 are bound to significantly affect the results. Second, incomplete and shallow genomes tend to present false negatives when mined for many symbionts. In addition, the origin of the DNA used for the genome sequencing will affect any conclusion on presence/absence or abundance of the symbionts detected and those undetected. In our study, all the used genomes came from DNA extracted from legs, therefore there is a methodical hard bias against gut fauna for example. Third, this kind of exploratory analyses of genomic material does not inform about the nature of the interaction between the organisms found in the genomic mix. Furthermore, in the majority of cases, this method also does not inform on the origin of the organisms. This is especially important as sample contamination is a known problem since the appearance of the molecular sequencing techniques. Finally, this method is not suitable for quantification of the present organisms. Altogether, these limitations exemplify the exploratory nature of the approach we used in this study.

## Conclusion

As we expected, our method detects various symbiotic partners in several Erebidae moth species, including *Wolbachia* and the bacteriophage *WO* in four species, *Burkholderia* in one other species, and *Sodalis* and *Arsenophonus* simultaneously in two species. Although symbiotic associations of Lepidoptera with *Wolbachia* is likely, similar long-term associations between the three other symbionts and the Lepidoptera have yet to be described. Similarly, we detect DNA material from bracoviruses that are currently only described as mutualistic symbionts of Hymenoptera. The true nature of these associations requires further experimental investigation. The detection of bracovirus DNA could for example suggest ecological interactions between moths and parasitoids, and the ability of the formers to naturally resist parasitoid attack strategies. Altogether our study presents a method and produces material supporting testable hypotheses about the diversity and nature of symbiotic interactions in those particular Lepidoptera species. With the availability of open access metagenomics databases, this field promises extensive and exciting opportunities to explore potentially hidden symbiotic diversity.

## Material and Methods

### Genome Data

We used the data produced from the whole genome sequencing project of 47 Erebidae species (see ^46^). The sampling information is shown in Table 1. This selection includes genomes representing the main described subfamilies and major lineages within the Erebidae family. The DNA was extracted from one or two legs of the selected samples. Extractions took place in 2000s / over a decade ago, for the purpose of another study (see ^47^). It is important to keep in mind that the genome sequencing approach generating this dataset is not optimized to recover the symbiont diversity of these organisms, therefore the diversity is likely to be systematically underestimated.

### Metagenomic analysis

The raw reads were quality checked with FASTQC v0.11.8 ^48^. Reads containing ambiguous bases were removed from the dataset using Prinseq 0.20.4 ^49^. Reads were cleaned to remove low quality bases from the beginning (LEADING: 3) and end (TRAILING: 3) and reads less than 30 bp in length. The evaluation of read quality with a sliding window approach was done in Trimmomatic 0.38 ^50^. Quality was measured for sliding windows of 4 bp and had to be greater than PHRED 25 on average. Cleaned reads were assigned taxonomic labels with Kraken2 ^51^ and MetaPhlAn 2.0 ^52^. Kraken2 was run using a custom database, which contained the standard kraken database, the refseq viral, bacteria and plasmid databases and all available Lepidoptera genomes from genbank (Supplementary Table 1 contains a full list of taxa included), confidence threshold of 0.05, and a mpa style output. MetaPhIAn was run using the analysis type rel_ab_w_read_stats, which provides the relative abundance and an estimate of read numbers originating from each clade. We visually screened the result for each sample, focusing on seven genera of vertically transmitted bacterial symbionts (i.e., *Arsenophonus* sp., *Cardinium* sp., *Hamiltonella* sp., *Rickettsia* sp., *Sodalis* sp., *Spiroplasma* sp. and *Wolbachia* sp.), one group of fungal symbionts (Microsporidia), and three types of viral symbionts (i.e., *Wolbachia*-phage *WO*, ichnovirus and bracovirus). This represents a non-exhaustive list of the maternally inherited symbionts found in diverse insect hosts, but covers all of those that have already been characterized within Lepidoptera ^10^. We also checked on the presence of the gut bacteria *Burkholderia* sp., which are known to confer pesticide resistance to their host in the pest bean bug *Riportus pedestris* (e.g., ‘can degrade an organophosphate pesticide, fenitrothion) ^53^.

## Data availability

The genome data used in this study are deposited in the NCBI SRA under BioProject PRJNA702831. All data in the supplementary material, the tables and the results can be found and downloaded from the GitHub repository: github.com/Hamidhrg/ErebidSymbionts.

## Acknowledgements

HRG received funding from the European Union’s Horizon 2020 research and innovation program under the Marie Sklodowska-Curie grant agreement No. 6422141. AD was funded by Marie Sklodowska-Curie Individual fellowship (#790531, Home Sweet Home). The authors acknowledge support from the National Genomics Infrastructure in Genomics Production Stockholm funded by Science for Life Laboratory, the Knut and Alice Wallenberg Foundation and the Swedish Research Council, and SNIC/Uppsala Multidisciplinary Center for Advanced Computational Science for assistance with massively parallel sequencing and access to the UPPMAX computational infrastructure. The authors highly value equity, diversity and inclusion in science. We would like to acknowledge the international character of our team, which significantly contributed to the completion and quality of the present study. It includes researchers from different countries, backgrounds and career stages.

## Author Contributions

H.R.G. and V.T. conceived the presented idea. H.R.G. carried out the experiments and wrote the manuscript with input from all authors. V.T. and A.D. designed the computational framework and analysed the data. All authors discussed the results and commented on the manuscript.

## References

1. Douglas, A. E. Multiorganismal Insects: Diversity and Function of Resident Microorganisms. Annual Review of Entomology 60, 17–34 (2015).

2. Dillon, R. J. & Dillon, V. M. THE GUT BACTERIA OF INSECTS: Nonpathogenic Interactions. Annual Review of Entomology 49, 71–92 (2004).

3. Duplouy, A., Hursts, G. D. D., O’neill, S. L. & Charlat, S. Rapid spread of male-killing Wolbachia in the butterfly Hypolimnas bolina. Journal of Evolutionary Biology 23, 231–235 (2010).

4. Altizer, S. M., Oberhauser, K. S. & Brower, L. P. Associations between host migration and the prevalence of a protozoan parasite in natural populations of adult monarch butterflies. Ecological Entomology 25, 125–139 (2000).

5. Jiggins, Hurst, Dolman & Majerus. High-prevalence male-killing Wolbachia in the butterfly Acraea encedana. Journal of Evolutionary Biology 13, 495–501 (2000).

6. Xu, P., Liu, Y., Graham, R. I., Wilson, K. & Wu, K. Densovirus Is a Mutualistic Symbiont of a Global Crop Pest (Helicoverpa armigera) and Protects against a Baculovirus and Bt Biopesticide. PLoS Pathogens 10, e1004490 (2014).

7. Bapatla, K. G., Singh, A., Yeddula, S. & Patil, R. H. Annotation of gut bacterial taxonomic and functional diversity in Spodoptera litura and Spilosoma obliqua. Journal of Basic Microbiology 58, 217–226 (2018).

8. Chen, F. et al. Effects of Wolbachia on mitochondrial DNA variation in populations of Athetis lepigone (Lepidoptera: Noctuidae) in China. Mitochondrial DNA Part A 28, 826–834 (2017).

9. van Nieukerken, E. J. et al. Order Lepidoptera Linnaeus, 1758. In: Zhang, Z.-Q. (Ed.) Animal biodiversity: An outline of higher-level classification and survey of taxonomic richness. Zootaxa 1758, 212–221 (2011).

10. Duplouy, A. & Hornett, E. A. Uncovering the hidden players in Lepidoptera biology: the heritable microbial endosymbionts. PeerJ 6, e4629 (2018).

11. Werren, J. H., Windsor, D. & Guo, L. Distribution of Wolbachia among neotropical arthropods. Proceedings of the Royal Society of London. Series B: Biological Sciences 262, 197–204 (1995).

12. Salunke, B. K. et al. Determination of Wolbachia Diversity in Butterflies from Western Ghats, India, by a Multigene Approach. Applied and Environmental Microbiology 78, 4458–4467 (2012).

13. Duplouy, A. & Brattström, O. Wolbachia in the Genus Bicyclus: a Forgotten Player. Microbial Ecology 75, 255–263 (2018).

14. Jiggins, F. M., Bentley, J. K., Majerus, M. E. & Hurst, G. D. How many species are infected with Wolbachia□? Cryptic sex ratio distorters revealed to be common by intensive sampling. Proceedings of the Royal Society of London. Series B: Biological Sciences 268, 1123–1126 (2001).

15. Tagami, Y. & Miura, K. Distribution and prevalence of Wolbachia in Japanese populations of Lepidoptera. Insect Molecular Biology 13, 359–364 (2004).

16. Ilinsky, Y. & Kosterin, O. E. Molecular diversity of Wolbachia in Lepidoptera: Prevalent allelic content and high recombination of MLST genes. Molecular Phylogenetics and Evolution 109, 164–179 (2017).

17. Sazama, E. J., Ouellette, S. P. & Wesner, J. S. Bacterial Endosymbionts Are Common Among, but not Necessarily Within, Insect Species. Environmental Entomology 48, 127–133 (2019).

18. Gavotte, L. et al. A Survey of the Bacteriophage WO in the Endosymbiotic Bacteria Wolbachia. Molecular Biology and Evolution 24, 427–435 (2006).

19. Wang, G. H. et al. Bacteriophage WO Can Mediate Horizontal Gene Transfer in Endosymbiotic Wolbachia Genomes. Frontiers in Microbiology 7, 1–16 (2016).

20. Wang, N., Jia, S., Xu, H., Liu, Y. & Huang, D. Multiple Horizontal Transfers of Bacteriophage WO and Host Wolbachia in Fig Wasps in a Closed Community. Frontiers in Microbiology 7, 1–10 (2016).

21. Tanaka, K., Furukawa, S., Nikoh, N., Sasaki, T. & Fukatsu, T. Complete WO Phage Sequences Reveal Their Dynamic Evolutionary Trajectories and Putative Functional Elements Required for Integration into the Wolbachia Genome. Applied and Environmental Microbiology 75, 5676–5686 (2009).

22. Kaushik, S., Sharma, K. K., Ramani, R. & Lakhanpaul, S. Detection of Wolbachia Phage (WO) in Indian Lac Insect [Kerria lacca (Kerr.)] and Its Implications. Indian Journal of Microbiology 59, 237–240 (2019).

23. LePage, D. P. et al. Prophage WO genes recapitulate and enhance Wolbachia-induced cytoplasmic incompatibility. Nature 543, 243–247 (2017).

24. Shropshire, J. D., On, J., Layton, E. M., Zhou, H. & Bordenstein, S. R. One prophage WO gene rescues cytoplasmic incompatibility in Drosophila melanogaster. Proceedings of the National Academy of Sciences 115, 4987 (2018).

25. Bordenstein, S. R. & Bordenstein, S. R. Eukaryotic association module in phage WO genomes from Wolbachia. Nature Communications 7, 13155 (2016).

26. Kent, B. N. & Bordenstein, S. R. Phage WO of Wolbachia: lambda of the endosymbiont world. Trends in Microbiology 18, 173–181 (2010).

27. Dale, C., Young, S. A., Haydon, D. T. & Welburn, S. C. The insect endosymbiont Sodalis glossinidius utilizes a type III secretion system for cell invasion. Proceedings of the National Academy of Sciences 98, 1883–1888 (2001).

28. Boyd, B. M. et al. Two Bacterial Genera, Sodalis and Rickettsia, Associated with the Seal Louse Proechinophthirus fluctus (Phthiraptera: Anoplura). Applied and Environmental Microbiology 82, 3185–3197 (2016).

29. Fukatsu, T. et al. Bacterial Endosymbiont of the Slender Pigeon Louse, Columbicola columbae, Allied to Endosymbionts of Grain Weevils and Tsetse Flies. Applied and Environmental Microbiology 73, 6660–6668 (2007).

30. Šochová, E., Husník, F., Nováková, E., Halajian, A. & Hypša, V. Arsenophonus and Sodalis replacements shape evolution of symbiosis in louse flies. PeerJ 5, e4099 (2017).

31. Burke, G. R., Normark, B. B., Favret, C. & Moran, N. A. Evolution and Diversity of Facultative Symbionts from the Aphid Subfamily Lachninae. Applied and Environmental Microbiology 75, 5328–5335 (2009).

32. Santos-Garcia, D., Silva, F. J., Morin, S., Dettner, K. & Kuechler, S. M. The All-Rounder Sodalis: A New Bacteriome-Associated Endosymbiont of the Lygaeoid Bug Henestaris halophilus (Heteroptera: Henestarinae) and a Critical Examination of Its Evolution. Genome Biology and Evolution 9, 2893–2910 (2017).

33. Toju, H. & Fukatsu, T. Diversity and infection prevalence of endosymbionts in natural populations of the chestnut weevil: relevance of local climate and host plants. Molecular Ecology 20, 853–868 (2011).

34. Conord, C. et al. Long-Term Evolutionary Stability of Bacterial Endosymbiosis in Curculionoidea: Additional Evidence of Symbiont Replacement in the Dryophthoridae Family. Molecular Biology and Evolution 25, 859–868 (2008).

35. Kaiwa, N. et al. Bacterial Symbionts of the Giant Jewel Stinkbug Eucorysses grandis (Hemiptera: Scutelleridae). Zoological Science 28, 169–174 (2011).

36. Rubin, B. E. R., Sanders, J. G., Turner, K. M., Pierce, N. E. & Kocher, S. D. Social behaviour in bees influences the abundance of Sodalis (Enterobacteriaceae) symbionts. Royal Society Open Science 5, 180369 (2018).

37. Sameshima, S., Hasegawa, E., Kitade, O., Minaka, N. & Matsumoto, T. Phylogenetic Comparison of Endosymbionts with Their Host Ants Based on Molecular Evidence. Zoological Science 16, 993–1000 (1999).

38. Oishi, S., Moriyama, M., Koga, R. & Fukatsu, T. Morphogenesis and development of midgut symbiotic organ of the stinkbug Plautia stali (Hemiptera: Pentatomidae). Zoological Letters 5, 16 (2019).

39. Mason, C. J. & Raffa, K. F. Acquisition and Structuring of Midgut Bacterial Communities in Gypsy Moth (Lepidoptera: Erebidae) Larvae. Environmental Entomology 43, 595–604 (2014).

40. Khojandi, N., Haselkorn, T. S., Eschbach, M. N., Naser, R. A. & DiSalvo, S. Intracellular Burkholderia Symbionts induce extracellular secondary infections; driving diverse host outcomes that vary by genotype and environment. ISME Journal 13, 2068–2081 (2019).

41. Itoh, H. et al. Host–symbiont specificity determined by microbe–microbe competition in an insect gut. Proceedings of the National Academy of Sciences of the United States of America 116, 22673–22682 (2019).

42. Ohbayashi, T., Itoh, H., Lachat, J., Kikuchi, Y. & Mergaert, P. Burkholderia gut symbionts associated with European and Japanese populations of the dock bug Coreus marginatus (Coreoidea: Coreidae). Microbes and Environments 34, 219–222 (2019).

43. Itoh, H. et al. Evidence of environmental and vertical transmission of Burkholderia symbionts in the oriental chinch bug, Cavelerius saccharivorus (Heteroptera: Blissidae). Applied and Environmental Microbiology 80, 5974–5983 (2014).

44. Kikuchi, Y. et al. Symbiont-mediated insecticide resistance. Proceedings of the National Academy of Sciences of the United States of America 109, 8618–8622 (2012).

45. Louis, F. et al. The Bracovirus Genome of the Parasitoid Wasp Cotesia congregata Is Amplified within 13 Replication Units, Including Sequences Not Packaged in the Particles. Journal of Virology 87, 9649–9660 (2013).

46. Ghanavi, H. R., Twort, V., Hartman, T. J., Zahiri, R. & Wahlberg, N. The (non) accuracy of mitochondrial genomes for family level phylogenetics: the case of erebid moths (Lepidoptera; Erebidae). bioRxiv (2021) doi:10.1101/2021.07.14.452330.

47. Zahiri, R. et al. Molecular phylogenetics of Erebidae (Lepidoptera, Noctuoidea). Systematic Entomology 37, 102–124 (2012).

48. Andrews, S. FastQC: A Quality Control Tool for High Throughput Sequence Data. http://www.bioinformatics.babraham.ac.uk/projects/fastqc/ (2010).

49. Schmieder, R. & Edwards, R. Quality control and preprocessing of metagenomic datasets. Bioinformatics (2011) doi:10.1093/bioinformatics/btr026.

50. Bolger, A. M., Lohse, M. & Usadel, B. Trimmomatic: A flexible trimmer for Illumina sequence data. Bioinformatics (2014) doi:10.1093/bioinformatics/btu170.

51. Wood, D. E. & Salzberg, S. L. Kraken: ultrafast metagenomic sequence classification using exact alignments. Genome Biology 15, R46 (2014).

52. Segata, N. et al. Metagenomic microbial community profiling using unique clade-specific marker genes. Nature Methods 9, 811–814 (2012).

53. Kikuchi, Y. & Yumoto, I. Efficient Colonization of the Bean Bug Riptortus pedestris by an Environmentally Transmitted Burkholderia Symbiont. Applied and Environmental Microbiology 79, 2088–2091 (2013).

